# Restoration legacy, landscape context and elevation shape frugivory in a tropical landscape

**DOI:** 10.64898/2025.12.18.694937

**Authors:** Pedro Luna, Juan E. Guevara-Andino, Betzabet Obando-Tello, Leonela E. Dávila-Narváez, H. Martin Schaefer, Nico Blüthgen

## Abstract

Assisted and passive natural restoration are widely applied strategies for forest restoration, yet their focus on tree recovery makes their effectiveness in restoring broader biodiversity unclear. Assuming that tree recruitment will rebuild whole ecosystems overlooks other taxa and their interactions, and thus underestimates key components of biodiversity like biotic interactions. To address this gap, we assessed two complementary questions: (i) how frugivory reestablishes and varies among assisted restoration and natural regeneration areas, and (ii) how local conditions (fruit availability, elevation, and time since restoration) and landscape context (forest cover and fragmentation) influence frugivory beyond restoration strategies. This was done by implementing a dummy fruit experiment in a tropical landscape in southeast Ecuador that considered two fruit sizes. Dummy fruit handling did not differ between assisted and naturally regenerated areas, indicating that neither restoration strategy was superior in promoting frugivory. Instead, landscape context played a central role. Fruit handling increased with old-growth forest cover and elevation, and declined with increasing forest fragmentation, highlighting the importance of habitat amount and connectivity for interaction recovery. Fruit size further shaped frugivory patterns, particularly in restored areas, where larger fruits were handled more frequently. Overall, our results show that restoration outcomes for frugivory depend less on restoration strategy alone and more on landscape structure and environmental context. By demonstrating the utility of artificial fruits as a rapid and practical tool to assess interaction recovery, this study highlights the need to move beyond vegetation-based metrics and explicitly incorporate biotic interactions into the evaluation of forest restoration success.

## Introduction

In light of the UN Decade on Ecosystem Restoration (2021–2030; UNEP & FAO, 2021), which calls for practical, evidence-based restoration of both species and ecosystem functions, it is vital to assess the performance of different restoration strategies. In this sense, restoring tropical rainforests through effective restoration practices will be essential for safeguarding global biodiversity, as these habitats are increasingly threatened by deforestation, habitat loss, and climate change (Dirzo and Raven, 2003; Pimm et al. 2014). Two common strategies are employed to mitigate these impacts: assisted natural restoration (active restoration) and passive natural restoration (passive restoration) (Holl and Aide, 2011; Chazdon et al. 2021). Although both approaches aim to reverse human-induced degradation and facilitate ecosystem recovery, they differ in their objectives, timelines, and levels of intervention. Each strategy has shown success in restoring forest cover and species diversity (Crouzeilles et al. 2017; Meli et al. 2017; Reid et al. 2018), yet evidence suggests that their outcomes are context-dependent, influenced by various confounding environmental and social factors (Holl and Aide 2011; Chazdon et al. 2021). Moreover, many restoration efforts have traditionally focused on recovering tree species diversity, often overlooking the recovery of ecological processes like biotic interactions (Selwyn et al. 2023; SER, 2021). To better evaluate the success of restoration, it is therefore necessary to assess how different strategies contribute to the recovery of ecosystem functions driven by interacting species. This understanding is critical for guiding restoration actions and meeting global goals in a rapidly changing world.

Evidence on whether assisted or passive natural restoration is more effective for restoring forest cover and biodiversity has yielded mixed results. While some studies indicate that assisted restoration leads to greater diversity recovery (Shoo et al. 2016; Holl et al. 2017; Hua et al. 2022), global meta-analyses indicate that passive natural restoration is more effective (Crouzeilles et al. 2017; Meli et al. 2017). Passive natural restoration is often considered a more efficient approach, as it is the most cost-effective and naturally regenerating forests tend to provide a higher return on investment in terms of biodiversity recovery (Crouzeilles et al. 2017). This conclusion primarily arises from the lower cost of passive natural restoration compared to active restoration, which can be significantly more expensive (Catterall and Harrison 2006). However, when assessing whether assisted or natural regeneration is the more effective strategy, it is crucial to consider the local context (e.g., historical land use, ecosystem resilience, and landscape factors), as these factors strongly influence regeneration success and the interpretation of study results, as evidenced by research supporting assisted regeneration (Shoo et al. 2016; Holl et al. 2017; Hua et al. 2022). Another aspect to consider is that meta-analyses favoring passive natural restoration may be subject to bias and require careful interpretation. For instance, Reid et al. (2018) identified a positive selection bias in meta-analyses: studies on natural regeneration were conducted in more forested regions, inherently favoring biodiversity measures in those areas. In contrast, assisted restoration was studied under more variable conditions, ranging from highly disturbed habitats to secondary forests, often including landscapes where natural recovery is hindered or stalled by low surrounding forest cover and past land use (Herrera et al. 2011; Laurance et al. 2011a; Reid et al. 2018). For instance, invasive plant species like grasses and ferns can hinder the establishment of dispersed seeds, delaying natural forest regeneration and promoting arrested recovery in these systems. In such degraded habitats, assisted regeneration may be essential to initiate the recovery process (Hartig and Beck, 2003; Roos et al. 2011; Levy-Tacher et al. 2015). Due to site-selection bias and the potential for stalled restoration, Reid et al. (2018) recommended focusing on paired comparisons within a region, ensuring that assisted and natural regeneration are assessed under the same ecological and environmental conditions. Indeed, studies that have conducted within-region comparisons found no consistent differences in restoration outcomes between natural regeneration and assisted restoration (Levy-Tacher et al. 2015; Shoo et al. 2016; Holl et al. 2017).

Frugivory is a key biotic interaction that drives forest regeneration, as frugivores are responsible for most seed dispersal in tropical ecosystems (Jordano, 2000). By dispersing seeds, frugivores facilitate seedling establishment and contribute to vegetation recovery in disturbed areas (Bello et al. 2024; Selwyn et al. 2023). However, many frugivores depend on forested areas for reproduction and foraging (Bonfim et al. 2021; Selwyn et al. 2023). Yet, the widespread loss of native forests has significantly reduced their original cover, threatening not only frugivore species (e.g., birds and mammals) but also their interactions with plants and the ecosystem services they sustain (Fahrig, 2003; Laurance et al. 2012; Fahrig et al. 2019). Evidence shows that in disturbed landscapes, frugivory is limited by forest loss and fragmentation, making restoration efforts crucial for the recovery and maintenance of these keystone interactions (Bonfim et al. 2021; Martínez-Penados et al. 2024). Despite the negative effects of forest loss and fragmentation on frugivores, their persistence and interactions can play a key role in supporting forest regeneration beyond remnant patches by enhancing seed dispersal and contributing to carbon sequestration in regenerating forests (Béllo Carvalho et al. 2023). Frugivore bird spill-over, for example, can occur from secondary and old-growth forest promoting the removal of fruits in disturbed habitats (Béllo Carvalho et al. 2023). However, frugivory dynamics can differ between conserved and disturbed habitats. Because, while some native frugivores facilitate regeneration by dispersing native seeds across both, habitat-generalist species are often linked to the spread of non-native seeds, potentially hindering restoration (Thierry et al. 2022). Moreover, although local frugivore species diversity can be similar across conserved and disturbed habitats, high functional turnover of frugivores and dispersed seeds has been observed which may increase the differences in fruit removal between conserved and degraded habitats (González-Varo et al. 2023). Thus, despite the potential of frugivores to support seed dispersal in regenerating forests, their identity and dynamics can vary with habitat condition. In this context, the extent to which habitat condition influences frugivory remains poorly understood, particularly regarding how different restoration strategies shape frugivore interactions within recovering areas.

Landscape ecology provides us with tools to understand how the composition (types and amounts of habitats) and configuration (spatial arrangement of habitats) of landscapes influence species and their interactions (Laurance et al. 2011; Haddad et al. 2015; Taubert et al. 2018; Arroyo-Rodríguez et al. 2020). In general, it is recognized that biodiversity is higher in habitats with high forest cover and low fragmentation which promote the dispersal of animals and plants leading to more diverse and healthy environments (Arroyo-Rodríguez et al. 2020; Corro et al. 2019; Bonfim et al. 2021). In the context of frugivory and restoration ecology, integrating landscape and restoration strategies may provide new insights into how restored ecosystems influence biotic interactions, enhancing our understanding of the factors that shape frugivory in disturbed areas. Additionally, elevation related environmental changes (i.e., decreases in temperature and air pressure, and increases in solar radiation with elevation) and temporal dynamics (i.e., succession and community assembly) may further shape biodiversity and ecosystem functions within restored landscapes (Körner, 2007; Walker et al. 2007; Crouzeilles et al. 2016), making them essential when evaluating restoration success.

In this study, we applied a landscape ecology approach and used old-growth forest as a reference to evaluate how frugivory interactions reestablish in a tropical cloud forest under restoration in southwest Ecuador. Specifically, we addressed two complementary questions: (i) how frugivory reestablishes and varies among assisted restoration and natural regeneration areas, and (ii) how local conditions and landscape context influence frugivory beyond restoration strategies. To address these questions, we conducted a fruit predation experiment using dummy fruits of two sizes (small vs. large). We hypothesized that assisted restoration areas would exhibit higher fruit handling rates than naturally regenerating areas, reflecting the targeted planting of fruiting tree species in the study area (Figure 1a). This expectation is also supported by previous evidence showing that frugivorous bird assemblages often recover more rapidly in assisted restoration areas, where tree planting accelerates the development of structural complexity and food resources (Morrison and Lindell 2011; Barros et al. 2022). In addition, across assisted and natural regeneration sites and old-growth forest, we expected smaller fruits to experience higher handling rates, since their size allows a broader subset of frugivores to access and consume them (Figure 1a; Morán-López et al. 2025). We then examined how local conditions (fruit availability, elevation, and time since restoration) and landscape context (forest cover and fragmentation) influenced fruit handling. We predicted that recently restored areas, particularly those characterized by lower forest cover and higher forest fragmentation, would exhibit reduced fruit handling rates compared to older restored and old-growth forest areas. Conversely, we predicted higher fruit handling rates in areas with older restoration, greater forest cover, and lower forest fragmentation (Figure 1b; Moral et al. 2007; Thierry et al. 2022; González-Varo et al. 2023). These expectations were based on evidence that landscapes with greater forest cover tend to support higher frugivore diversity (Laurance et al. 2011; Reid et al. 2014; Bonfim et al. 2021), and that reduced forest connectivity associated with forest loss and fragmentation can negatively affect the recovery of frugivory interactions (Landim et al. 2025). Fruit availability was also expected to influence frugivory, given that variation in resource abundance can affect fruit–frugivore interactions in disturbed landscapes (Herrera et al. 2011). Along the elevational gradient, we predict higher levels of fruit handling at lower elevations, where warmer and more stable climatic conditions typically support greater frugivore richness and abundance, potentially increasing encounter rates between frugivores and fruits (Ferger et al. 2016).

**Figure 1.**
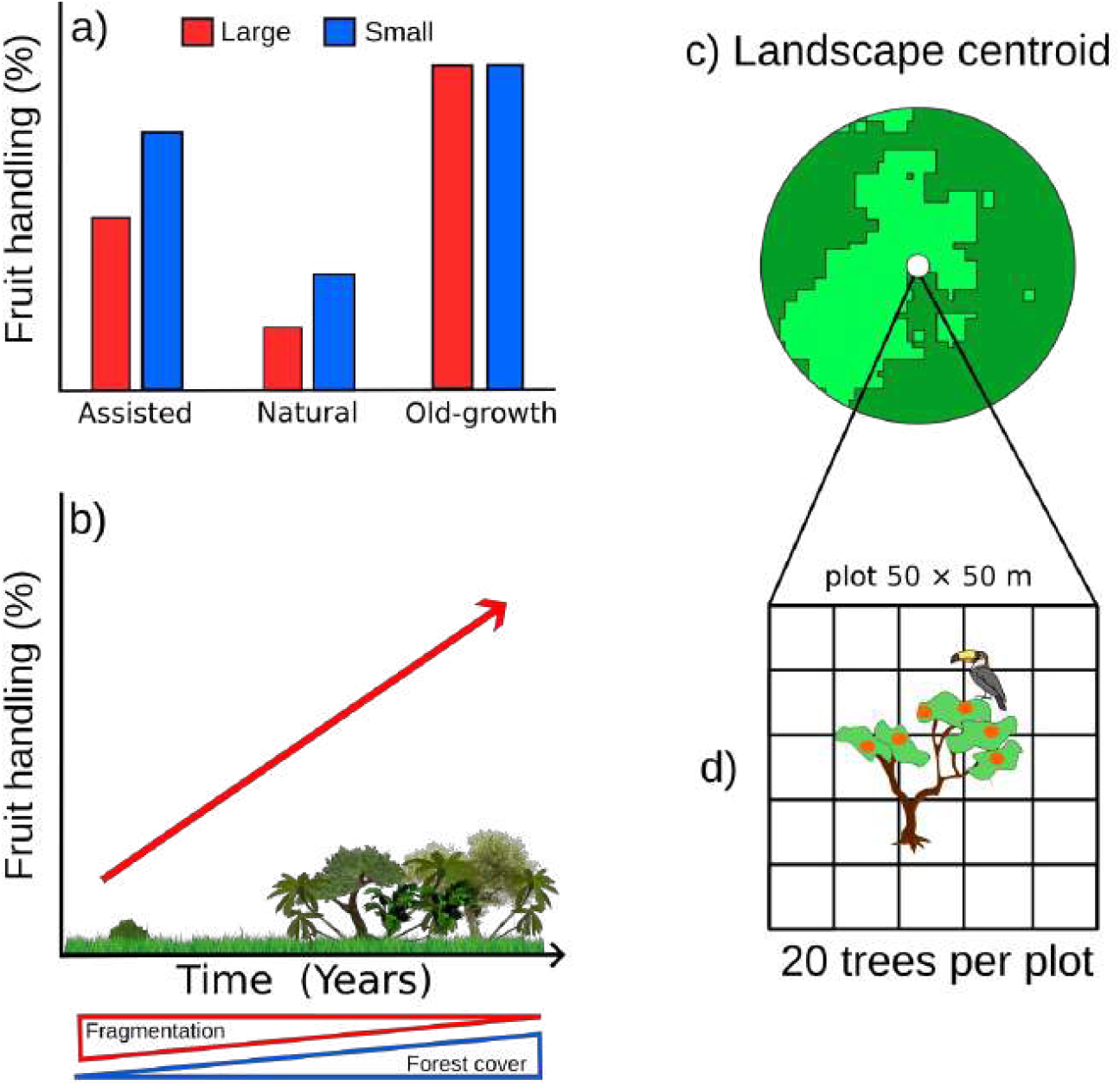
(a) Expected proportion of fruit handling events on dummy fruits across restoration strategies: assisted natural regeneration (assisted), passive natural regeneration (natural), and reference old-growth forest (old-growth). (b) Expected proportion of fruit handling events in relation to landscape composition (forest cover) and landscape configuration (fragmentation). (c) Example of the landscape buffer showing the centroid at which each sampling plot was located for the assessment of frugivory. (d) Schematic representation of the experimental design implemented at each landscape centroid (n = 21).

## Methods

### Study site

This study was conducted in the private reserve Buenaventura (Jocotoco Foundation), located in the province of El Oro in southeastern Ecuador (3°39’13” S, 79°46’4” W). The reserve spans 4,200 hectares and ranges in elevation from 450 to 2,200 meters above sea level. The habitat within Buenaventura reserve is influenced by the Choco biodiversity hotspot and it harbors some of the last remaining and certainly the largest tropical mountain cloud forest of the region, underscoring its high conservation value. However, these forest remnants are surrounded by a highly degraded landscape primarily due to intensive livestock farming. This land use has transformed the natural landscape into a matrix of old-growth forest, restored secondary forest and grasslands dominated by Merker grass (*Pennisetum purpureum*, Poaceae) and the invasive bracken fern (*Pteridium aquilinum*, Dennstaedtiaceae). Merker grass, an invasive species from Africa, grows rapidly up to 4 meters in height within three months and out-competes native plants, posing significant challenges for reforestation efforts by impeding the growth of newly planted trees. Bracken fern is a competitive invasive species that produces a dense frond canopy that reduces the amount of light reaching the herbaceous layer of vegetation. It also develops extensive rhizome system that can act as a barrier which prevents the establishment of native plant species. Both Merker grass and Bracken fern represent a challenge for the success of restoration efforts, as they are very hard to control and their presence stalls natural restoration (Hartig and Beck, 2003; Roos et al. 2011; Levy-Tacher et al. 2015 et al.).

Restoration strategies in the Buenaventura Reserve are categorized into assisted natural regeneration and passive natural regeneration and have been underway since the reserve’s establishment in 1999 (approximately 26 years). Assisted natural regeneration involves planting native tree species (up to 18 different species focusing on fast growing and fruiting trees like *Miconia* sp., *Cecropia* sp., *Ochroma* sp. and *Ficus* sp.) in areas formerly dominated by Merker grass, Bracken fern or a mixture of both. Reforestation teams maintain tree plantations by mowing the surrounding grass and ferns every three months for 3 years (until tree seedlings reach heights that surpass grass and fern cover). Planted trees contribute to forest cover recovery by attracting frugivores and combating the invasive weeds by creating shaded areas that inhibit its spread.

Passive natural regeneration involves protecting areas previously used for livestock to allow them to recover autonomously with minimal human intervention beyond ensuring protection. In naturally regenerated areas, isolated trees are present and are expected to perform similar ecological roles as those in actively restored areas, though their abundance is generally lower. Old-growth forests areas are present within the reserve; however, they are highly fragmented, especially near areas previously used for livestock and those currently undergoing restoration.

In addition to the restoration context, the Buenaventura Reserve spans a marked elevational gradient associated with changes in temperature, ranging from mean 21.6 °C (± 1.72 SD) at 484 m a.s.l. to 17.54 °C (± 2.58 SD) at 1711 m a.s.l. Temperature data were recorded using 10 HOBO data loggers deployed across the elevational gradient of the reserve.

### Landscape selection and categorization

In the Buenaventura Reserve, we selected 21 sampling points (mean distance ± SD = 651.15 ± 448.37 m) across three distinct land management types: assisted regeneration, passive natural regeneration, and old-growth forest, with seven sampling points per category. Each sampling point was treated as the centroid of a landscape buffer used to characterize the surrounding forest cover composition, which included varying proportions of grasslands, secondary forest, and old-growth forest (Figure 1c). For each landscape centroid, we recorded elevation and the age of the sampling point (hereafter referred to as time), defined either as the time elapsed since restoration began in restored areas or as the estimated stand age in old-growth forest. Following the selection of sampling points, we quantified forest cover in the surrounding landscape using a supervised classification based on a tree height raster with a spatial resolution of 10 m (Lang et al. 2023). Field-based measurements of tree height were used to establish threshold values for forest classification, allowing us to distinguish forested from non-forested areas according to the observed range of canopy heights within the study plots. Based on this classification, the raster was used to define three forest cover categories, which were validated using tree height data from 50 field reference points. These categories were defined as: (i) old-growth forest (tree height range: 27–44 m); ii) secondary forest (20 to 26 m); iii) regenerated areas and grass or fern dominated areas (0 to 19 m). Using the classified raster, we quantified landscape composition (forest cover) and landscape configuration (edge density) for each landscape across six spatial extents, defined by concentric buffer radii ranging from 100 to 600 m at 100 m intervals (Figure S1 & S2). Forest cover and edge density were calculated from the classified raster using the R packages “multilandr” (Huais 2024) and “terra” (Hijmans 2020). For the purposes of this study, forest cover refers specifically to the proportion of old-growth forest within each buffer and was represented as a binary variable (old-growth forest = 1; non-forest = 0), where the non-forest category included secondary forest, restored areas, and grasslands. Edge density represents the amount of edge between old-growth forest and the surrounding matrix (secondary forest, restored areas, grassland and fern-dominated areas) per unit area within each buffer, with higher values indicating greater forest fragmentation. All landscape classification and map processing were conducted in QGIS version 3.22.4 and R version 4.4.1 (R Core Team 2025).

### Frugivory experiment

To obtain a consistent proxy of frugivory across the different land management types, we conducted an experiment using artificial fruits of two sizes (small, 10 mm diameter, and large, 20 mm diameter) made from non-toxic red plasticine (Pelikan ®). For this, at the centroid of each landscape, we established a 50 × 50 m plot, where we selected 20 young, fruitless trees (ranging from 1 to 1.8 meters in height) and placed 6 dummy fruits of one size (either small or large) on each tree (Figure 1c). In total, we deployed 120 dummy fruits per sampling point (i.e., buffer centroid) (60 small and 60 large), with 2,520 dummy fruits across all landscapes (1,260 small and 1,260 large) (Figure 1d).

After placing the dummy fruits, we left them exposed for 72 hours. After this time, we returned to each sampling point and recorded whether each fruit had been handled (i.e., showed evidence of interaction) or not by potential frugivores. Fruit handling was recorded as a binary response (handled = 1, not handled = 0), allowing us to calculate the proportion of handled fruits of each size within each landscape. We also documented the type of marks on each fruit, identifying whether they were made by birds, mammals, or arthropods (Figure S3). Fruit monitoring was conducted directly on focal trees.

When a fruit was no longer present on a focal tree, the surrounding area was searched to identify the type of mark associated with fruit handling. If the fruit could not be located, it was classified as handled, but the type of mark was not recorded.

After recording the markings, we collected the remaining fruits. Note that plasticine dummy fruits offer a standardized, durable, and non-toxic tool for controlled ecological experiments, enabling a consistent measure of frugivore events.

### Fruit availability

In order to have a measure of fruit availability at each site we made a sampling by counting and estimating all the fruits that were observed within the 50 × 50 m plot. This was made by the same observer in all landscapes using binoculars and a hand tally counter. This was done to assess if fruit availability influences attack rates on dummy fruits.

### Data analysis

#### Effect of restoration strategy

As a first step in understanding how fruit handling varies across the Buenaventura Reserve, we examined the variation in the types of markings observed on dummy fruits. Specifically, we assessed whether markings caused by birds, arthropods, or mammals differed between fruit sizes (small and large) and across restoration strategies. We used a generalized linear model (GLM) with binomial error distribution with link *logit* with the proportion of handled fruits as response variable with type of marking (bird, arthropod, mammal or other), fruit size (small and large) and restoration strategy (assisted, natural restoration and old-growth forest) as explanatory factors. Fruits with unidentified markings were excluded from this analysis.

After identifying the main frugivores handling the dummy fruits, we assessed how fruit handling varied among restoration strategies while accounting for fruit size. To do so, we excluded all frugivory events attributed to arthropods, as well as markings of unidentified origin, to avoid potential confounding effects. This decision was based on our aim to specifically assess frugivory by vertebrates, namely birds and mammals. We then fitted a generalized linear model with a binomial error distribution and *logit* link function, using the proportion of handled fruits as the response variable and fruit size (small vs. large) and restoration strategy (assisted regeneration, passive natural regeneration, and old-growth forest) as explanatory factors. We used likelihood ratio tests to assess the significance of predictors and tested for differences among categories using post hoc pairwise comparisons with false discovery rate correction. Analyses were conducted using the “car” library (Fox and Weisberg 2011) for generalized linear models and “emmeans” (Lenth 2025) for post hoc tests in base R version 4.4.1 (R Core Team, 2025).

#### Effect of local conditions and landscape context

To assess the effect of local conditions (fruit availability, elevation, and time since restoration) and landscape context (forest cover and edge density) on fruit handling we first identified the scale-of-effect for the landscape descriptors. This consisted of determining the buffer radius at which the proportion of handled fruits best responded to landscape context. We did this by generating a series of models in which forest cover, edge density and landscape heterogeneity measured at six buffer radii (100, 200, 300, 400, 500 and 600 m) were tested as predictors of the proportion of handled fruits. For each model we extracted the AICc value and selected the buffer radius with the lowest AICs as the scale of effect for that metric. After identifying the optimal buffer size for forest cover (100 m buffer) and edge density (400 m buffer) for each response variable (Figure S4), we fitted a global model. The global model included elevation, time since restoration, fruit availability, forest cover, and edge density as predictors, with the proportion of handled fruits as the response variable. The landscape descriptors used in this model corresponded to the buffer radii previously identified as having the strongest support in the scale-of-effect analysis. We assessed multicollinearity among predictors using the variance inflation factor (VIF). This analysis indicated low multicollinearity among predictors (VIF < 3), and therefore no variables were excluded. We then applied a multimodel inference approach based on information criteria, using AICc (Akaike Information Criterion corrected for small sample sizes). Differences among models were quantified using ΔAICc, calculated as the difference between the AICc value of each model and the lowest AICc value, with the best-supported model having ΔAICc = 0. After identifying the best model, we used likelihood ratio tests to assess the significance of the retained predictors. The best model retained elevation, forest cover and edge density. All predictors were standardized previous to model selection and the models were analyzed using a Binomial error distribution with a *logit* link function. Model selection was performed using the “MuMIn” (Bartón 2010) library and generalized linear models were done using “car” (Fox and Weisberg 2011) and “stats” library in base R version 4.4.1 (R Core Team, 2025).

In addition to our analysis and in order to understand Buenaventura landscape, we described matrix composition around sampling points, by quantifying the proportional cover of major land cover classes (old-growth forest, secondary forest, and regenerated areas and grass or fern dominated areas) within circular buffers of 100 m and 400 m radius around each landscape centroid. These buffer sizes correspond to the spatial scales at which forest cover and edge density showed the strongest support in the scale-of-effect analysis. Land-cover proportions were summarized descriptively and visualized using stacked bar plots for the matrix surrounding each sampling point (Figure S2). We also tested for correlations among covariates using Spearman correlation test among local conditions (fruit availability, elevation, and time since restoration) and landscape factors (forest cover and edge density).

Additionally, we assessed the role of spatial autocorrelation in the proportion of attacked fruits by using Moran’s I and correlograms. For this, we first grouped all pairwise plot distances into 12 bins of equal sample size (centroids: 410 m to 10 500 m). Distance bins were built so that each bin contained approximately the same number of pairwise distances. This ensures equal statistical power across bins, even though the distance ranges vary in order to achieve equal counts. For each bin, we ran 999 permutations to obtain the observed Moran’s I, its expected value under randomness, 95 % confidence limits, and a p-value. This was done with the package “letsR” (Vilela and Villalobos, 2015) in R version 4.4.1 (R Core Team, 2025).

## Results

### Effect of restoration strategy

A total of 803 fruits (31% of all dummy fruits) were attacked, including 373 small fruits (15%) and 430 large fruits (16%). Among the attacked fruits, 669 (83%) were marked by birds (361 large, 308 small), 30 (3.7%) by mammals (23 large, 7 small) and 51 (6.3%) by arthropods (37 large, 14 small). Additionally, 53 fruits (6.6%) were either lost or had unidentified markings (9 large, 44 small) (Figure S3). Bird markings were the most frequent type of attack across both fruit sizes (χ^2^=1349, df = 3, P = 0.0001). Larger fruits were more frequently attacked than smaller ones across all marking types (χ^2^=5.23, df = 1, P = 0.02) (Figure 2a). Marking type either made by birds, mammals or arthropods were similar among assisted, natural regeneration areas and old-growth forest (χ^2^ = 2.87, df = 2, P = 0.23) (Figure 2b).

**Figure 2.**
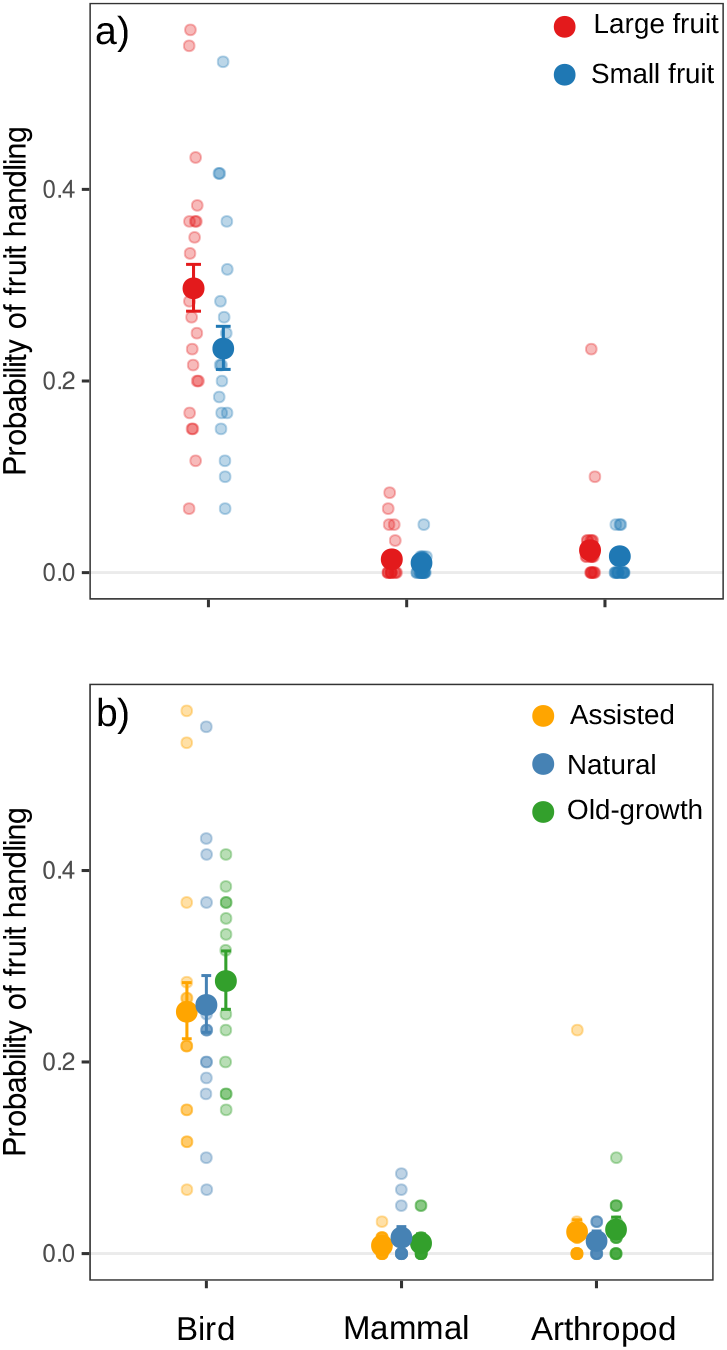
Predicted probability of fruit handling a) by fruit size and type of mark and b) by restoration strategy and type of mark. Circles show predicted probabilities of fruit handling, and error bars indicate 95% confidence intervals. Scattered points show observed proportions. Unidentified markings were excluded from this analysis.

When fruit size is not considered, we found that fruit handling did not differ among restoration strategies or between restored areas and old-growth forest (χ^2^ = 2.51, df = 2, P = 0.28). However, fruit size had an effect on fruit handling (χ^2^ = 9.45, df = 1, P = 0.002), and its interaction with restoration strategy revealed contrasting patterns between restored and old-growth forest areas (χ^2^ = 7.82, df = 2, P = 0.02; Figure 3). In this sense, we observed that large fruits were handled more frequently than small fruits in assisted restoration (predicted mean probability [95% CI]: large = 0.29 [0.25, 0.34]; small = 0.23 [0.19, 0.27]) and natural regeneration areas (large = 0.33 [0.29, 0.38]; small = 0.22 [0.18, 0.26]). In contrast, in old-growth forest, fruits of both sizes were handled at similar rates (large = 0.29 [0.25, 0.34]; small = 0.3 [0.26, 0.35]), resulting in overall higher fruit handling probabilities in forest sites when both fruit sizes are considered together. Importantly, the main differences between old-growth forest and restored areas were not reflected in absolute differences in proportions, but rather in contrasting size-dependent trends, which may indicate different frugivory dynamics across restoration contexts.

**Figure 3.**
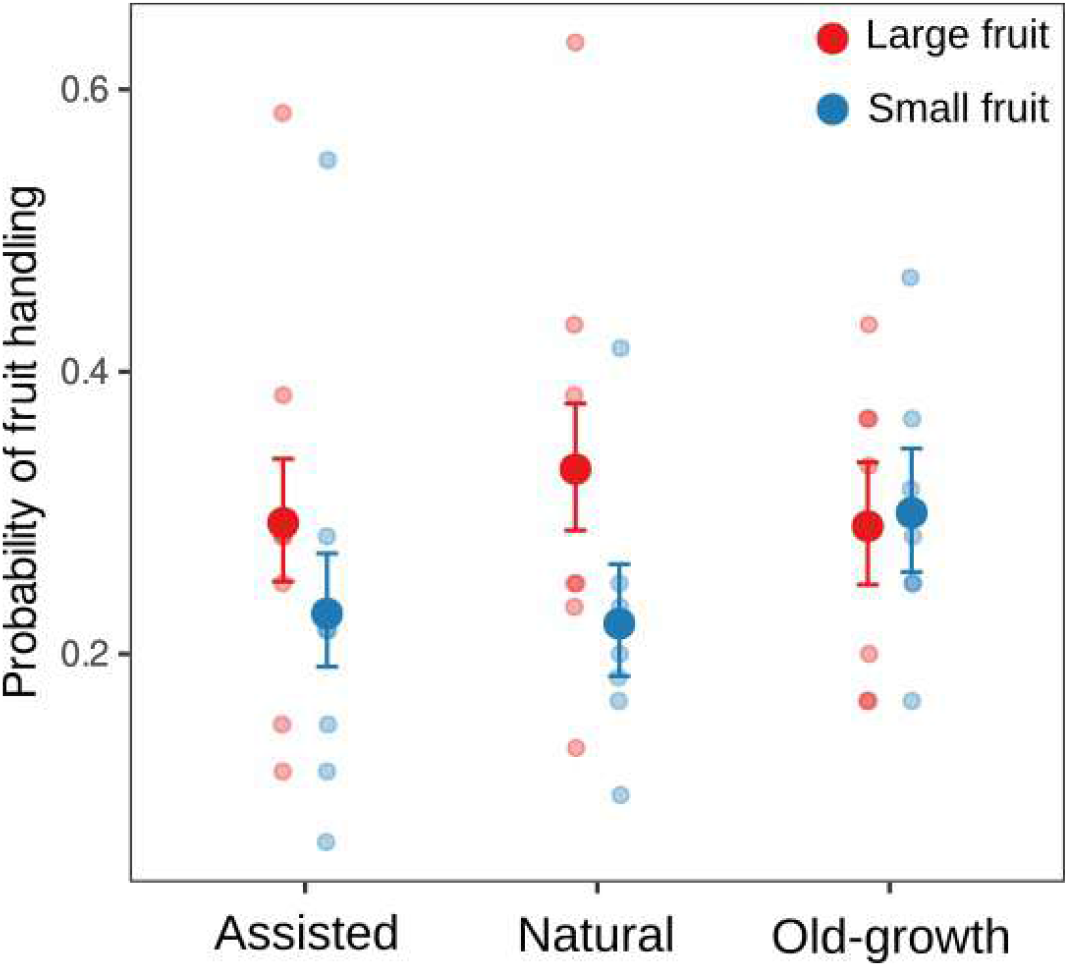
Probability of fruit handling across restoration strategies (assisted, natural regeneration, and old-growth forest) for small and large dummy fruits. Circles show predicted probabilities of fruit handling, and error bars indicate 95% confidence intervals. Scattered points show observed proportions. Arthropod and unidentified markings were excluded from this analysis.

### Effect of local conditions and landscape context

When assessing which factors best explained fruit handling, we found that the probability of fruit handling increased with elevation and forest cover, while it decreased with edge density (Table 1; Figure 4). This indicates that frugivory is more frequent in areas with greater forest cover and at higher elevations, whereas increased landscape fragmentation has a negative effect on fruit handling.

**Table 1.** Deviance table showing the contribution of each predictor to the probability of fruit handling based on likelihood ratio tests. Reported coefficients (β) represent standardized effect sizes. LR χ^2^ values and associated P-values indicate the significance of each predictor. Bold values denote statistically significant effects (P < 0.05).

| Factor | $\beta$ | LR $\chi^2$ | df | P |
| --- | --- | --- | --- | --- |
| Elevation | 0.21 | 17.61 | 1 | < **0.0001** |
| Forest cover | 0.24 | 23.34 | 1 | < **0.0001** |
| Edge density | -0.17 | 12.52 | 1 | < **0.001** |

**Figure 4.**
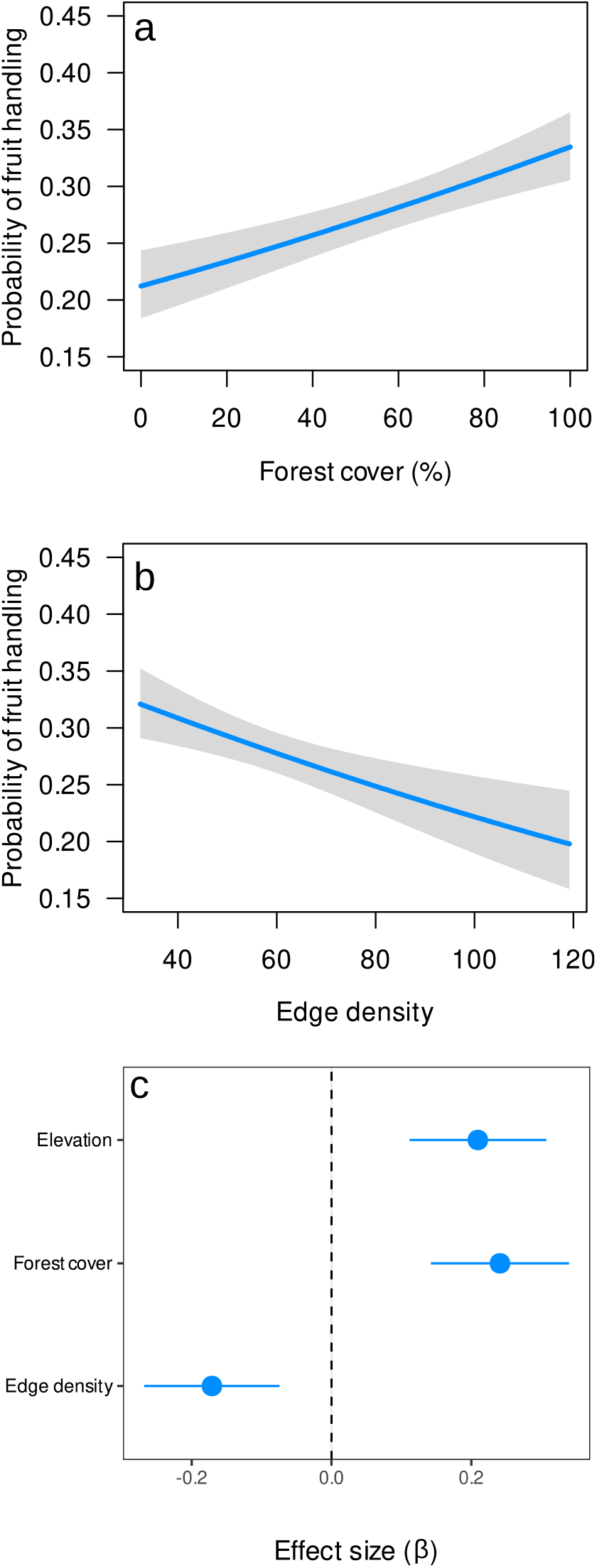
Effect of local conditions and landscape context on the probability of fruit handling. Panels show predicted probabilities for (a) forest cover, (b) edge density, and (c) effect size. Panel (d) shows standardized effect sizes (β ± 95% confidence intervals) for the predictors retained in the final model.

Matrix composition varied among landscapes and across spatial scales (Figure S1 & S2). At the local scale (100 m buffer), mean old-growth forest cover across landscapes was 61.5% (± 38.9 SD), while secondary forest and grassland accounted for 23.7% (± 25.9) and 14.8% (± 26.8), respectively. At the broader scale (400 m buffer), mean old-growth forest cover was 58.1% (± 26.6), with secondary forest and grassland contributing 27.7% (± 16.5) and 14.3% (± 13.2), respectively. Correlations among local conditions and landscape factors indicated that areas with greater forest cover are also the older ones (ρ = 0.76, P < 0.001). In addition, sampling points at higher elevations tended to be younger (ρ = −0.36, P = 0.01) and to have lower forest cover in their surroundings (ρ = −0.48, P < 0.001) (Figure S5).

The Moran’s I spatial autocorrelation analysis did not show significant clustering on the frequency of attacks on fruits across different distances (up to 6 km). At longer distances (around 8 kilometers), landscapes showed signs of dissimilarity (Moran’s I = -0.35, P < 0.001). These results suggest a shift from no spatial autocorrelation at shorter distances to dispersion at greater distances (Figure S6).

## Discussion

Identifying where and how animals interact with plants remains a major challenge in restoration ecology, as much of the theoretical and empirical work has focused on species gains and losses rather than on the recovery or disruption of biotic interactions. In this study, using a controlled experiment, we found evidence that frugivory is recovering in both assisted and naturally restored areas; however, we found no support for one restoration strategy being superior to the other. Instead, our results indicate that landscape context plays a central role in shaping the recovery of frugivory. Forest cover emerged as a key driver, suggesting that standing forest remnants act as reservoirs of frugivores that can spill over into surrounding restored areas. In contrast, forest fragmentation had a negative effect on fruit handling, highlighting the importance of landscape connectivity for the reestablishment of plant-animal interactions. Beyond restoration strategy and landscape structure, elevation also emerged as a predictor of fruit handling, with higher probabilities observed at higher elevations, underscoring the influence of environmental gradients on the recovery of frugivory. Our results also showed that our methodology has applied relevance, as we show that a simple and practical experimental approach using artificial fruits can be used to monitor the recovery of biotic interactions in restored landscapes. This methodology provides a rapid way to go beyond assessing vegetation recovery and to evaluate how biotic interactions, such as frugivory, are reestablishing. In this sense, our results reinforce the idea that restoring tree cover alone may not be sufficient, as even after 25 years of restoration efforts, frugivory rates still differ between restored areas and old-growth forest, with standing forests being key for the dispersal and recovery of animal and plant communities.

When accounting for the effects of restoration strategies on fruit handling, we observed that it occurred at similar rates in assisted and natural restoration areas, indicating that active reforestation is not necessarily more effective than passive natural regeneration in restoring frugivory. This result was more evident when considering fruit size, as both assisted and natural restoration areas showed similar trends, with large fruits being handled more frequently than small fruits. In contrast, old-growth forest showed a different pattern, with fruits of both sizes being handled at similar rates. Thus, the main difference between restored areas and old-growth forest was not the overall level of fruit handling, but rather the way fruit handling varied with fruit size. This suggests that, although frugivory is being recovered in restored areas, its dynamics differ from those observed in reference sites. The higher handling of larger fruits in restored areas may be related to the spillover of larger-bodied frugivorous birds from nearby forest remnants into restoration sites. This interpretation is supported by our observations that birds were the primary frugivores and by the fact that fruit size constrains which frugivores can interact with fruits, with larger birds generally capable of handling larger fruits. In our study site, spillover effects are likely, as several large frugivorous bird species, including toucans (*Ramphastos ambiguus, R. brevis* and *Pteroglossus torquatus*), guans (*Ortalis erythroptera, Penelope purpurascens* and *Chamaepetes goudotti*), and trogons (*Trogon collaris* and *Pharomachrus auriceps*), were frequently observed foraging near forest edges and within restoration sites. Moreover, assisted and natural restoration areas were, on average, located less than 170 m from old-growth forest, distances that are well within the movement ranges of large frugivorous birds such as toucans, which can move over distances of up to 2 km (Holbrook & Loiselle 2007; Holbrook 2011). These short distances, combined with the high mobility of frugivores, likely facilitate frequent movements across habitat types, promoting fruit handling in restored areas. This interpretation is consistent with previous work showing that birds tend to move more frequently from forested habitats into degraded areas than in the opposite direction, thereby sustaining fruit removal and seed dispersal in disturbed landscapes (Béllo Carvalho et al. 2023).

Despite the similar frugivory patterns observed between assisted and natural restoration, the contribution of assisted restoration to frugivory recovery in the Buenaventura Reserve should not be overlooked. In several areas, natural regeneration is constrained by invasive grasses and ferns, which can stall successional processes. Without human intervention, many of these sites would likely not reach conditions that support frugivory dynamics comparable to those observed in naturally regenerating areas (Hartig & Beck 2003; Roos et al. 2011; Levy-Tacher et al. 2015). In this context, assisted restoration can provide suitable habitat for frugivores in areas where natural regeneration is limited, potentially facilitating secondary growth by enabling seed consumption and dispersal similar to that occurring in naturally regenerating sites. In the case of the frugivory trends observed in old-growth forest we need also to consider that its difference with restoration areas may also be influenced by vertical stratification of frugivores. Because our experiment focused on understory frugivory, interactions by larger birds and mammals that forage primarily in the canopy may have been underrepresented (Thiel et al. 2021). Reduced handling of small fruits in restored areas and reduced handling of large fruits in forest may have implications for seed dispersal, potentially limiting dispersal opportunities for small-fruited early-successional species and large-fruited late-successional species, respectively (Galetti & Dirzo 2013; Bello et al. 2015; Galetti et al. 2013). Finally, fruit traits may further contribute to the observed patterns. The red fruits used in our experiment are expected to be particularly attractive to visually oriented frugivores such as birds. In restoration areas, where tree density and canopy cover are lower, large and conspicuous fruits may be more visible and thus more likely to attract a subset of larger frugivores, whereas smaller fruits may be less detectable. In contrast, many mammals rely more strongly on olfactory cues when locating fruits, which may partly explain the low rates of fruit handling by mammals observed in our study. These patterns are broadly consistent with the dispersal syndrome hypothesis, which predicts that fruit traits such as color, size, and scent are associated with the sensory capacities of different frugivore guilds (Valenta & Nevo 2020).

Forest cover emerged as the main driver of frugivory, indicating that areas surrounded by old-growth forest are those where fruit handling is more frequent. This result was expected, as old-growth forests typically exhibit higher structural complexity and resource availability, providing suitable habitat for a greater diversity of frugivorous birds and plants which increases the opportunities for plant–animal interactions to occur (Guariguata & Ostertag 2001; Chazdon 2014; Bonfim et al. 2021). However, we also found that in Buenaventura reserve old-growth forest patches are highly fragmented and embedded within a matrix of secondary forest, young restoration areas, and grasslands which ultimately exert negative impacts on frugivory. Although frugivory may occur in habitats other than old-growth forest, forested areas remain a high priority for the reestablishment and conservation of frugivory, as they function as reservoirs from which frugivores, particularly birds, can spill over into surrounding restored habitats (Herrera et al. 2011; Reid et al. 2014; Bello et al. 2024). Moreover, the negative effect of fragmentation further highlights that not only the amount of forest cover, but also the spatial continuity of high-quality habitat, is critical for sustaining frugivory. In this context, restoration strategies in the Buenaventura Reserve could benefit from explicitly promoting connectivity among forest patches, for example through targeted tree planting aimed at reducing local forest fragmentation. Although secondary forests represent a significant amount of the habitats within the reserve and may play an important role in supporting frugivores (Acevedo-Charry & Aide 2019), they recover slowly and may not fully replace old-growth forest, as they frequently lack forest specialists and key functional groups (Letcher et al. 2015; Rozendaal et al. 2019). Consequently, conserving frugivory in this landscape likely requires a dual strategy that protects remaining old-growth forest fragments while promoting the expansion and maturation of secondary forests, as has been proposed for maintaining tree species diversity (Rozendaal et al. 2019). Time since restoration is also playing an important role, as older sites tended to be associated with higher forest cover. This pattern suggests that the passage of time contributes to the gradual recovery of frugivore–plant interactions. As restored areas mature, they develop greater structural complexity, including taller canopies and more developed understory layers (Guariguata & Ostertag 2001; Lebrija-Trejos et al. 2008; Escobar et al. 2025), as well as increased fruit availability, all of which can attract a broader range of frugivores. Taken together, our results indicate that maintaining frugivory in Buenaventura reserve depends on safeguarding old-growth forest remnants, reducing fragmentation, and allowing restored and secondary forests sufficient time to mature and develop the structural and functional attributes necessary to support frugivore communities.

In lowlands, species richness and abundance tend to be higher due to warm, stable climates (McCain & Grytnes 2010; Rahbek et al. 2019), which generally promotes more interactions (Roslin et al. 2017; Hargreaves et al. 2019). Contrary to this expectation, fruit handling increased with elevation across our 400–1 700 m a.s.l. gradient, an effect that held regardless of forest cover or restoration strategy. We measured fruit availability along this gradient but found no significant correlation with elevation, indicating that increased frugivory at greater heights is not simply explained by differences in fruit abundance. Therefore, we could argue that the higher frugivory rates may reflect higher frugivore abundance, as interaction frequencies are largely driven by species abundance (Krishna et al. 2008). Although we did not directly sample frugivore communities, our results are consistent with previous studies reporting non-linear elevational patterns in frugivory. For example, Terborgh (1977) found that frugivorous bird diversity in the Andes peaked between 1 400 and 1 600 m, likely due to the local abundance of fruiting species in families such as Ericaceae, Rubiaceae, and Melastomataceae. The observed mismatch between fruit availability and attack rates suggests that other factors, such as frugivore community composition, foraging behavior, or microclimatic conditions, may explain the pattern. To better understand these dynamics, future research should combine elevation surveys of frugivores with detailed fruit trait data and microclimate measurements to evaluate their combined influence on frugivory across the elevation gradient studied.

Overall, our results show that the recovery of frugivory in a restored landscape depends less on the restoration strategy *per se* and more on the surrounding landscape context and local environmental conditions. Both assisted and naturally regenerated areas supported comparable levels of fruit handling, indicating that frugivory can recover under different restoration pathways. However, forest cover emerged as key determinants of this recovery, emphasizing the importance of conserving old-growth forest remnants and reducing fragmentation to sustain frugivore–plant interactions. Differences in fruit handling dynamics between restored areas and old-growth forest further suggest that, although frugivory is reestablishing, its functional patterns may differ from those in reference sites. Together, these findings reinforce the need to move beyond vegetation-based indicators of restoration success and to explicitly consider landscape structure, local context, and biotic interactions when evaluating the functional outcomes of tropical forest restoration.

## Acknowledgements

We thank Fundación de Conservación Jocotoco for granting permits and providing logistical support during fieldwork. We also thank Leovigildo Cabrera (son), Baldomiro Becerra and Leovigildo Cabrera (father) for their invaluable assistance in the field. PL is grateful to all Fundación Jocotoco Buenaventura park rangers, whose support was vital to this project. PL thanks Claudia Suarez for her support during the development of this project ily.

## Statements & Declarations

## Author Contributions

Original idea was developed by Pedro Luna, with input from Juan E. Guevara-Andino, H. Martin Schaefer and Nico Blüthgen. Pedro Luna, Betzabet Obando-Tello and Leonela E. Dávila-Narváez carried out field work. Pedro Luna carried out map manipulation and data analysis. The first draft of the manuscript was written by Pedro Luna and all authors commented on previous versions of the manuscript. All authors read and approved the final manuscript.

## Data availability

The datasets and maps generated during and/or analysed during the current study are available from the corresponding author on request.

