## Supplementary material for "Restoration legacy, landscape context and elevation shape frugivory in a tropical landscape"

**Submitted to Biodiversity and Conservation**

**Figure S1.** Land cover categories surrounding the sampling points. Circles denote concentric buffers ranging from 100 m to 600 m. Colors show the different land cover categories grassland and young restored areas (red), secondary forest cover (yellow) and old-growth forest (green). Codes above the buffer show the identity of each landscape centroid with A denoting a sampling point located in assisted restoration areas, N natural regeneration areas and F old-growth forests.

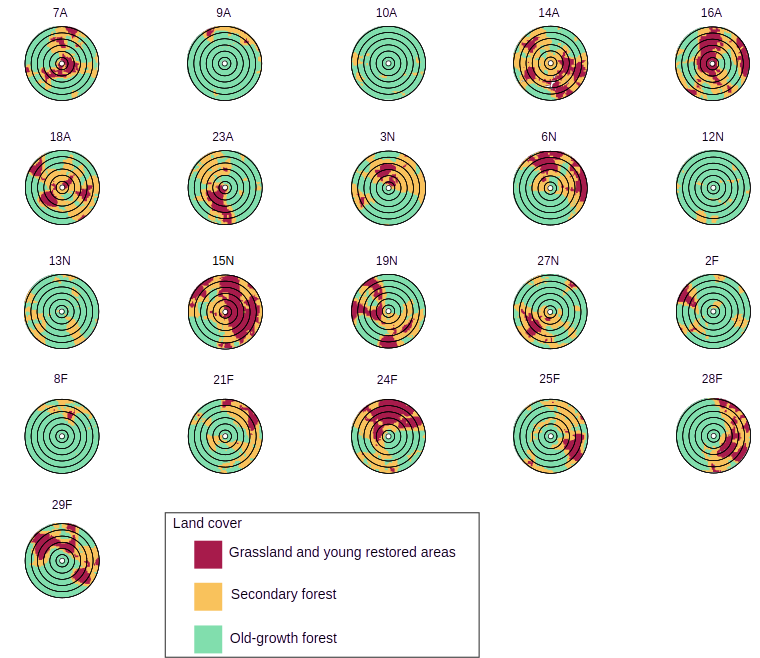

**Figure S2.** Proportional matrix composition of major land cover classes: grasslands and young regenerated areas (red), secondary forest (yellow) and old-growth forest (green), within circular buffers of 100 m and 400 m radius around each landscape centroids. Codes in the x-axis represent the identity of each landscape centroid with A denoting a sampling point located in assisted restoration areas, N natural regeneration areas and F old-growth forests.

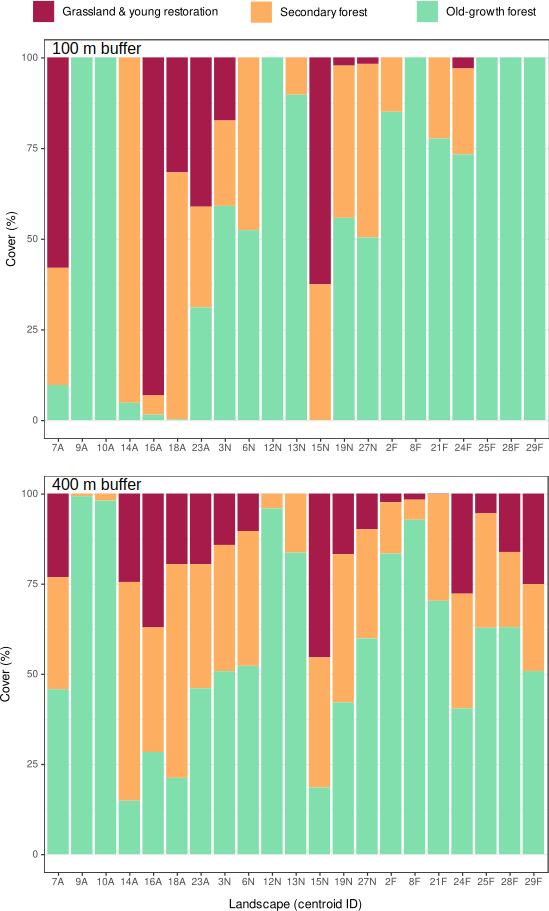

**Figure S3.** Examples of markings made by bird, mammals, arthropod and other (unidentified markings) on dummy fruits.

**Figure S4.** Scale-of-effect analysis showing variation in AICc values across buffer radii (100–600 m) for models explaining fruit handling as a function of forest cover and edge density. Lower AICc values indicate better model fit, and the buffer radius at which AICc is minimized represents the scale of effect for each landscape metric.

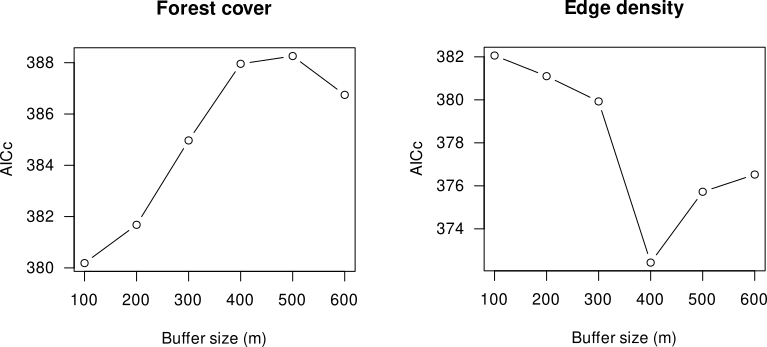

**Figure S5.** Correlation between local conditions (fruit availability, elevation, and time since restoration) and landscape context (forest cover and edge density). T = time since restoration; FC = forest cover (%); E= Elevation m a.s.l.; FA= fruit availability (%); and ED = edge density. Correlation test were made with Spearman’s rank correlation test. Only significative (P < 0.05) correlations are shown.

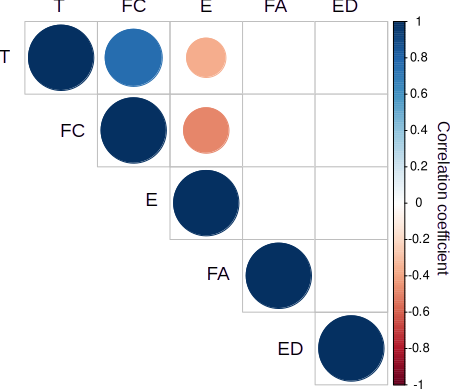

**Figure S6.** Correlogram showing the variation of spatial autocorrelation on the proportion of attacked fruits across the studied landscapes. Points denote observed Moran’s I values and bars their confidence intervals of 95%.

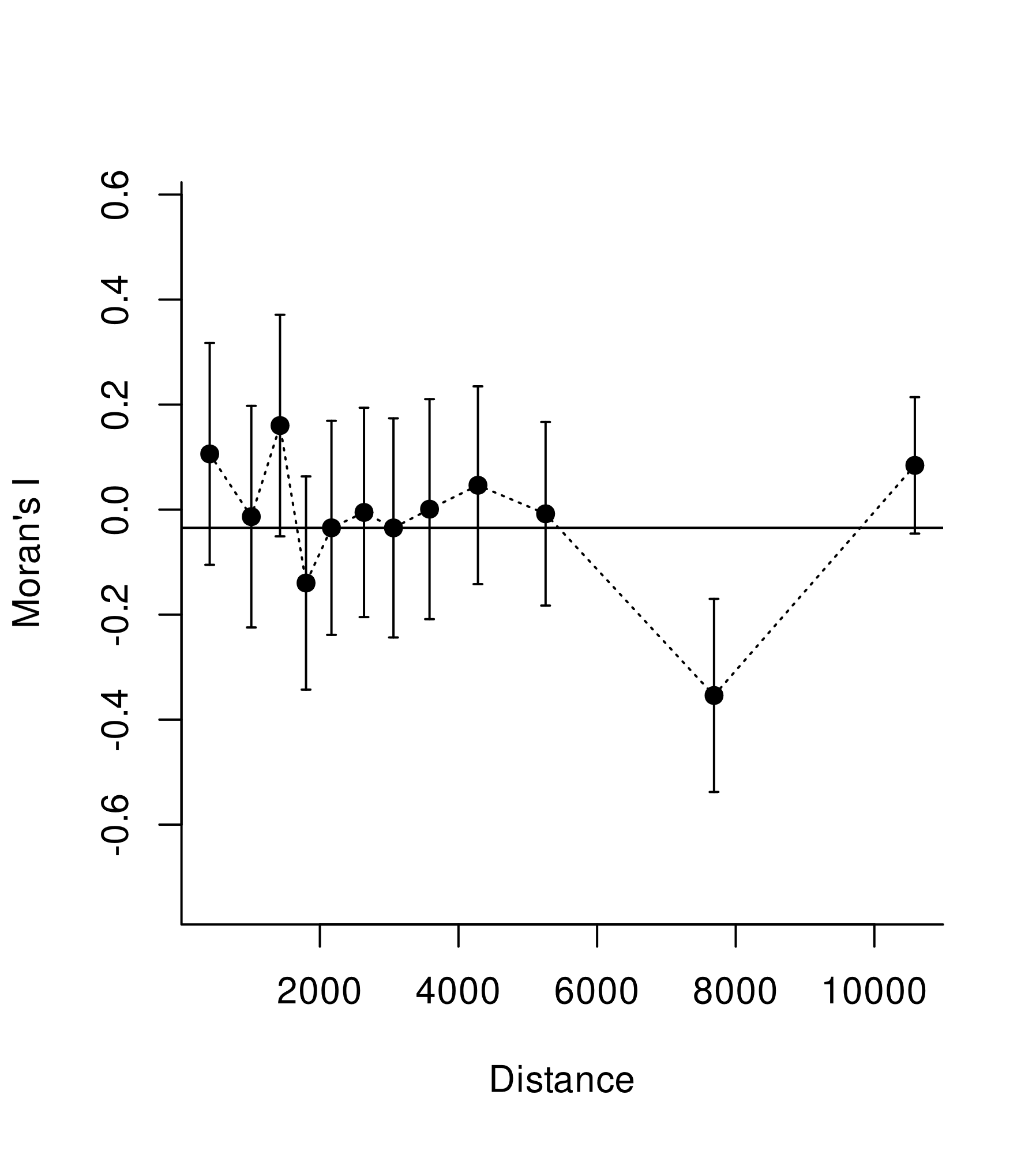
